# An experimental validation test of ecological coexistence theory to forecast extinction under rising temperatures

**DOI:** 10.1101/2024.02.22.581553

**Authors:** J. Christopher D. Terry

**Affiliations:** Department of Biology, University of Oxford, 11A Mansfield Road, Oxford, OX1 3SZ

**Keywords:** coexistence, climate change, Drosophila, Bayesian, ecological forecasting

## Abstract

Interactions between species pose considerable challenges for forecasting the response of ecological communities to global changes. Coexistence theory could address this challenge by defining the conditions species can or cannot persist alongside competitors. However, although coexistence theory is increasingly deployed for projections, these frameworks have rarely been subjected to critical multigenerational validation tests. Here, using a highly replicated mesocosm experiment, I directly test if the modern coexistence theory approach can predict time-to-extirpation in the face of rising temperatures within the context of competition from a heat-tolerant species. Competition hastened expiration and the modelled point of coexistence breakdown overlapped with mean observations under both steady temperature increases and with additional environmental stochasticity. That said, although the theory identified the interactive effect between the stressors, predictive precision was low even in this simplified system. Nonetheless, these results support the careful use of coexistence modelling for forecasts and understanding drivers of change.

## Introduction

Species are already shifting their distributions across the world as environmental conditions rapidly change (Chen *et al*. 2011; Lenoir *et al*. 2020). Forecasting how the identities of species resident in ecosystems will change is both of profound applied importance (Pecl *et al*. 2017) and a key test of our ecological understanding of community assembly. Beyond direct physiological responses, the potentially large influence of interspecific competition in determining whether a species can persist under a different environment has been demonstrated in both mesocosm (Davis *et al*. 1998; Legault *et al*. 2020) and field studies (Alexander *et al*. 2015; Descombes *et al*. 2020; Freeman *et al*. 2022; Suttle *et al*. 2007). Parsing this complexity of multi-species dynamics to make forecasts is a considerable challenge given the range of possible processes, but theoretical ecology has developed streamlined frameworks defining precise conditions for when species are, or are not, expected to coexist (Armitage & Jones 2020; Barabás *et al*. 2018; Chesson 2000; Ellner *et al*. 2019; Ranjan *et al*. 2024).

Modern coexistence theory focuses on a widely applicable currency: invasion growth rate (Chesson 2000; Grainger *et al*. 2019), the per-capita population growth rate from low densities into a stationary community of established competitors. Assuming no positive density-dependence (‘Allee effects’ Courchamp *et al*. 1999), a positive invasion growth rate is taken to imply that a species is able to persist on the basis that it could always recover from low densities. In a two-species system, when both species fulfil the mathematical criteria (Barabás *et al*. 2018), they are claimed to be able to coexist in a locality. This approach is frequently framed as an assessment of whether average fitness differences between pairs of species that would otherwise drive exclusion can be overcome by stabilising niche differences (Chesson 2000).

There have been increasing efforts to deploy this extensive body of coexistence theory for more applied purposes (Alexander *et al*. 2016; Aoyama *et al*. 2022; Hallett *et al*. 2023; Letten *et al*. 2021; Ocampo-Ariza *et al*. 2018) with dozens of applications to empirical systems (reviewed in Buche *et al*. 2022; Terry & Armitage 2024). As part of this, coexistence theory has been proposed as a tool to understand how environmental change will impact communities in the future, largely based on parametrising models from short-term experiments under different environmental conditions (Adler *et al*. 2012; Alexander *et al*. 2015, 2018; Bimler *et al*. 2018; Mordecai 2013; Van Dyke *et al*. 2022; Wainwright *et al*. 2019). Where key model parameters are found to be environmentally dependent, the future environments in which species can (or cannot) coexist can be inferred. An ultimate hope is to be able to make trait or phylogeny-based predictions (Godoy *et al*. 2014; Kraft *et al*. 2015; Van Dyke *et al*. 2022) of niche and fitness differences in different environments, potentially unlocking the capacity to forecast species responses to climatic changes at scale without individual parameterisation.

Since the initial development of ‘modern’ coexistence theory, it has been criticised from many angles - from the fundamental framing (Simha *et al*. 2022), through to the simplicity of how interspecific interactions are represented (Abrams 2022), and the challenge of accurately estimating the form of competition (Cervantes-Loreto *et al*. 2023). The underlying mathematical assumptions of standard modern coexistence theory are also widely recognised as being some distance from reality: criticisms include the focus on pairwise dynamics (Chesson 2018; Spaak & Schreiber 2023) and the potential misalignment of the spatial scales of observation and coexistence (Clark *et al*. 2019; Hart *et al*. 2017). Non-stationary environments generate transient dynamics that may be poorly represented by predictions derived from static observations (Chesson 2017). To calculate long-term growth rates the standard approach implicitly assumes fixed traits with infinite time and space horizons (Schreiber *et al*. 2023), as well as continuous population counts without the demographic stochasticity that discrete individuals at small population sizes bring (Pande *et al*. 2020). However, the extent to which these issues preclude the applicability of the framework to contemporary challenges remains unclear, as although it is applied to empirical systems with increasing frequency, its predictions have rarely been directly tested with multi-generational experiments. As such, the extent to which this framework is adequate (Getz *et al*. 2018) for ecological questions given natural complexities remains an unsettled question.

To assess the capacity for modern coexistence theory to forecast the time of extirpations (i.e. local extinctions) under changing environments, I conducted a highly replicated *Drosophila* mesocosm experiment where populations were tracked through discrete generations in small mesocosms (Figure 1). The controlled environment isolates several of the potentially troublesome simplifications by limiting the diversity and defining the spatial scale. This directly exposes to testing the assumptions of competition model sufficiency, infinite time and space horizons, as well as the potential impact of positive density dependence and adaptation.

**Figure 1.**
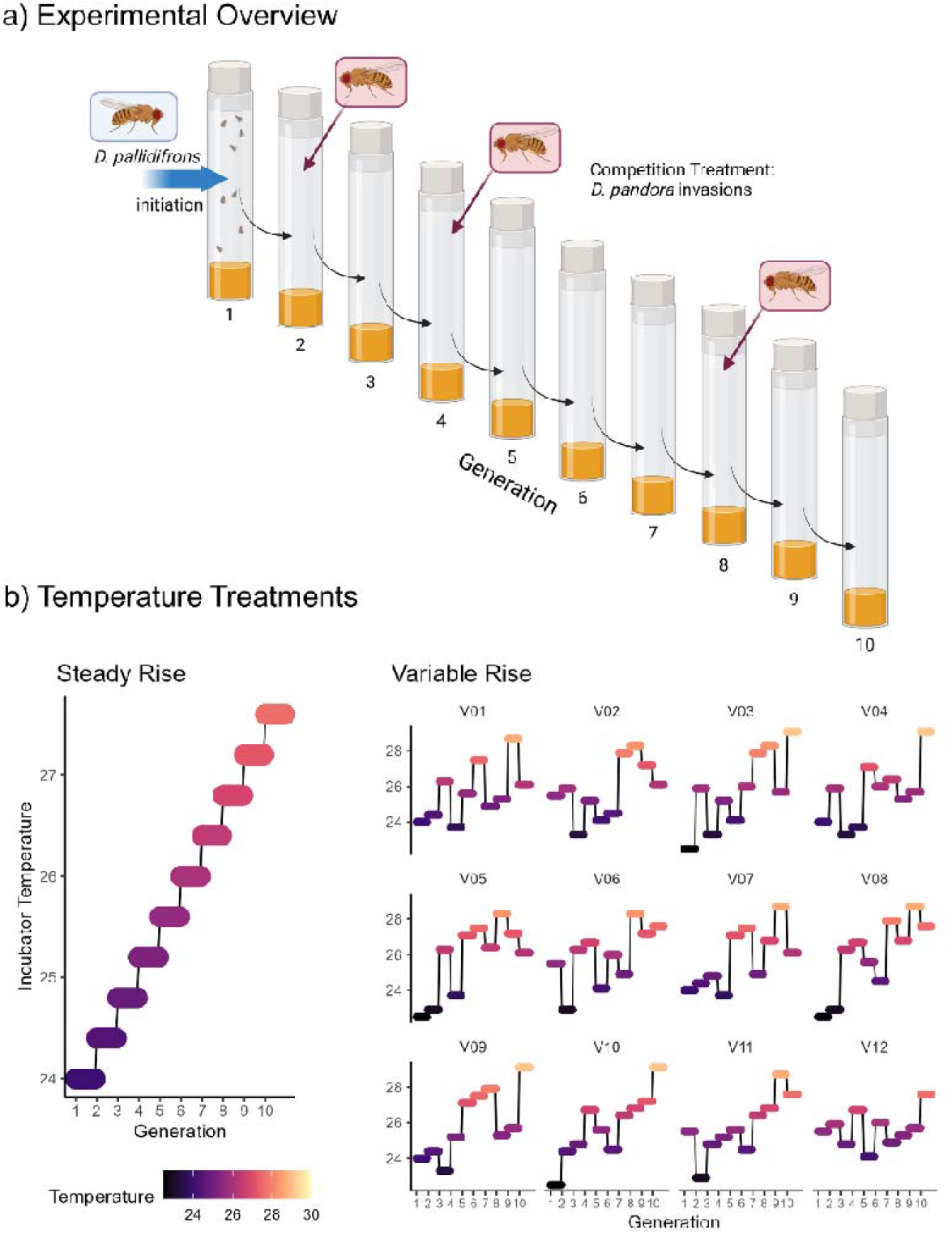
Outline of experiment and time-to-extinction results. a) Overall experiment schematic. After initiation of population vials with *D. pallidifrons*, the flies were allowed two days to lay eggs before removal and censusing. After a further 10 days, all emerged flies were transferred to a new vial for two days to start the next discrete generation. In the ‘competition’ treatment, small numbers of *D. pandora* were added alongside the founders of generation 2, 4 and 8. b) Temperature treatments. In the ‘steady rise’ scenario, the temperature increases 0.4°C at the start of each generation, while in the variable rise treatment, the temperature imposed on each population follows one of 12 trajectories (V01 – V12) that superimpose variability on top of the steady rise.

## Methods

I focussed on predicting the survival of populations of *Drosophila pallidifrons*, a highland-distributed species with a comparatively cool thermal optimum, in the context of competition from a lowland-distributed species *Drosophila pandora* with a comparatively warm thermal optimum. Previous empirical work using this community has demonstrated roles for both temperature and competition in determining species range boundaries (Chen & Lewis 2024; O’Brien *et al*. 2017).

### Species

The two species, the ‘resident’ *Drosophila pallidifrons* (*nasuta* species complex) and the ‘invader’ *Drosophila pandora* (*ananassae* subgroup, McEvey & Schiffer 2015), are both drawn from a well characterised montane Australian tropical rainforest community that shows distinct elevational community turnover (Hangartner *et al*. 2015; Jeffs *et al*. 2021; Kellermann *et al*. 2012). Laboratory populations had been maintained at Oxford University since 2018, originally drawn from multiple isofemale lines sampled from rainforest sites in northern Queensland, Australia (permit PWS2016-AU-002018, Australian Department of Energy and Environment). These cultures had been maintained for several years at large population sizes in a 25°C controlled temperature room. While some laboratory adaptation over this period cannot not be ruled out, preceding work using these populations had demonstrated that species had maintained distinct thermal physiologies (Chen & Lewis 2024).

### Experimental Procedures

Each generation was initiated by transferring the founder flies into a standard 25 mm diameter *Drosophila* vial with 5 ml of cornflour–sugar-yeast–agar *Drosophila* medium. The founders were allowed approximately 48 hours to lay eggs before being removed and frozen for later censusing. After 10 days incubation any emerged flies became the founders of the next generation in that line, giving a total tip-to-tip generation time of 12 days. This length was chosen to match the development time across the temperatures used in the experiment. During censusing, all individuals were identified by species, sexed and counted under a stereo microscope. Flies that had clearly died before freezing were not counted. Each generation 1 (G1) vial was initiated with 3 female and 2 male *D. pallidifrons* counted under light CO_2_ anaesthesia that had been raised in 180 ml Drosophila flasks under moderate density at 24°C. Populations were censused each generation and the experiment was run for 10 discrete generations (at which point 98.75% of *D. pallidifrons* populations were extinct). Sixty replicates were run for each of two crossed treatment combinations: monoculture or the intermittent introduction of small numbers of *D. pandora*, (Figure 1a), and steady temperature rise or with additional generational-scale thermal variability (either +1.5°C, -1.5°C or 0, with equal chance, Figure 1B).

In the ‘Steady’ rise temperature treatment, G1 was founded at a constant temperature of 24°C and the temperature was raised by a 0.4°C step at the start of each generation, implying a 4°C rise across the experiment. This was expected to shift the competitive balance from *D. pallidifrons* to *D. pandora*. For the ‘Variable’ rise treatment, 12 different variable temperature trajectories (Figure 1b) were generated by randomly assigning a third of the trajectories to each of lowered (−1.5), mean (=) or raised (+1.5) temperature conditions for each generation. Each trajectory had 5 replicates within each treatment. Temperature values were chosen to span the expected shift in coexistence, while also allowing the use of a fixed generation duration across the experiment (since colder temperatures delay development significantly). Experimental populations were maintained in three incubators of two equivalent models (Sanyo MIR-154 & Sanyo MIR-153) under a 12-12hr light-dark cycle. Temperature and humidity loggers indicated these incubators were able to successfully maintain a controlled environment throughout the experiment.

In the competition biotic treatment, *D. pandora* were intermittently added alongside the founders from the preceding generation. At the start of G2, approximately 15 *D. pandora* were added to each treatment vial. At the start of G4 and G8 4 female and 2 male *D. pandora* were added to all lines that still had a *D. pallidifrons* population. These additions were subtracted from the subsequent census to infer the reproductive rate in the previous generation.

### Population Data

In total 37,826 *D. pallidifrons* and 20,786 *D. pandora* flies were counted from 1,045 censuses with non-zero counts (Figure S1). Population dynamics were based just on female counts to better account for the reproductive capacity of small populations and since the sex ratio of both species skewed towards females at higher densities (Figure S2, likely due to slower emergence of males and increased male mortality). Across all temperatures, there were 1,223 *D. pallidifrons* and 592 *D. pandora* observed transitions from non-zero female populations. Because there were so few (6) *D. pallidifrons* data points from temperatures above 28°C, these were excluded to leave a total data set of 1,217 transitions. This allowed the lower bound of *T*_*max*_(see below) to be reduced without introducing numerical errors.

### Population Model Fitting

Models representing the generation-generation population dynamics of the two species were built following recently established best practices (Terry & Armitage 2024), including propagation of uncertainty in model parameters (Bowler *et al*. 2022) and consideration of alternative model forms (Cervantes-Loreto *et al*. 2023). A suite of phenomenological models were tested that systematically varied the model form used to represent the fundamental thermal performance curve, the reduction in fecundity caused by competition and the error distribution. These models were fit separately for each species.

All models were fit in a Bayesian framework using the *brms* version 2.19 (Bürkner 2018) R interface to STAN version 2.26.1 (Carpenter *et al*. 2017). Priors on the core growth rate function parameters were chosen to be weakly informative and bounded only to prevent numerical errors. Priors on the competition model parameters were based on existing expectations of the strength of competition (Chen & Lewis 2024; Terry *et al*. 2021) and to enforce competition rather than positive interactions. For each model, 4 chains were each run for 4000 total iterations (2000 used as a warm up) on standard settings. Model code in R (*brms*) and STAN is available in the code supplement. Runs were checked for convergence through the Gelman diagnostic 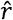. Posterior distributions and predictive plots are available in the code supplement. Model performance was compared using differences in ELPD-LOO (Vehtari *et al*. 2017), which assesses each model on its capacity to make predictions on unseen further data across the whole predictive distribution. Where models could not be clearly distinguished on the basis of predictive performance (with an ELPD difference significantly greater than 4) the simpler model was used.

Each model followed the overall structure:

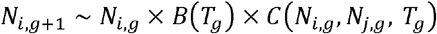

where *N* is the number of individual females and *T* the temperature indexed by the generation *g*, with *i* and *j* indicating focal and competitor species respectively. Based on existing expectations of the shape of thermal performance curves from this *Drosophila* community (Chen & Lewis 2024), four phenomenological asymmetric thermal performance curve models *B*(*T*_*t*_) were trialled, selected for having a moderate number of parameters and their distinctiveness (Kontopoulos *et al*. 2024): ‘Taylor-Sexton’, ‘Atkin’, ‘Simplified β-type’ and ‘Simplified Brière’. Only the Simplified Brière thermal performance function (Briere *et al*. 1999) was able to consistently converge:

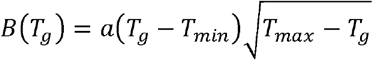

where *a, T*_*min*_ and *T*_*max*_ are parameters of the thermal performance function defining the low-density growth rate.

Eight phenomenological competition response models (Table S1) were tested for *C* (*N*_*i,t*_, *N*_*j,t*_, *T*_*t*_). These were based on generalisations of the widely used Beverton-Holt model and included temperature-dependent competition coefficients. Linear relationships (i.e. a Lotka-Volterra model) were also tested but converged poorly as they frequently generated negative expectations and were excluded from consideration. For both species, the selected competition kernel was the Beverton-Holt model of density-dependent competition with the competitive pressure exerted by *D. pallidifrons* negatively impacted by temperature:

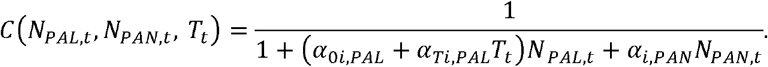

Where *α* terms are competition coefficients. The *α*_*ij*_ terms represent constant competition coefficients, whereas *α*_*Tij*_ is an intercept and *α*_*Tij*_ a slope term to linearly link a competition coefficient to temperature. Note that the number of intraspecific competitors of species was taken to be *max*, (0, *N* − 1) incorporating the assumption that individuals do not compete with themselves. Models including temperature dependent competition consistently performed better than models that did not (Table S3-4). For *D. pallidifrons*, the highest-ranked model included an additional exponent term, but this did not justify the additional model complexity (ΔELPD =6.3, SE = 2.5).

Six error models for count data were tested: Poisson, Negative Binomial, Zero-inflated Negative Binomial, Temperature-dependent-zero-inflated Negative Binomial, Zero-inflated Negative Binomial with competition dependent shape, and Zero-inflated Negative Binomial with temperature dependent shape (Table S2). The differences in error functions had a greater impact on ELPD-LOO than choice of competition model (Table S3-4). Error for *D. pallidifrons* was best represented by a zero-inflated negative binomial model, where the extent of zero inflation was temperature dependent (via a logit link function). For *D. pandora*, error was best represented by a negative binomial function where the shape parameter (ϕ) was competition dependent. See Table S5 and S6 for model parameter values and Figure S4 temperature dependence of emergent parameters.

### Predictions of coexistence at different temperatures

The selected population dynamic models were used to forecast the temperature at which *D. pallidifrons* would be extirpated (excluded by *D. pandora*) by three alternative approaches: 1) calculation of mutual invasibility using analytical modern coexistence theory via analysis of niche and fitness differences (‘classic’ modern coexistence theory), 2) Calculation of long-term growth rates from simulations following the method of Ellner et al. (2019), and 3) Direct replicate simulations of expected extinctions in the specified scenario.

### Predictions from analytic coexistence theory

Following the original specification of niche and fitness differences (Chesson 1990; Spaak *et al*. 2023) derived for the Beverton-Holt model in Godoy et al. (2014), I defined the fitness difference between the two species as:

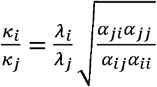

and the niche overlap as

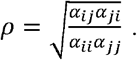

The temperature dependent zero-inflation was incorporated into the overall growth rate term *λ* by scaling the mean expectation of the number of offspring by the zero inflation: *λ* (*T*_*t*_) = *B* (*T*_*t*_) × (1 − *inv. logit* (*z*_0_ + *z*_*T*_ *T*_*t*_)). Predicted coexistence at a steady state of each temperature was inferred based on whether the key inequality was fulfilled (Chesson 2000): 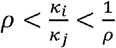. The full posterior parameter estimates of the best performing model was propagated through to give relative support for each of the four coexistence outcomes (exclusion of either species, coexistence or priority effects whereby both species can exclude the other) at each temperature.

### Prediction of coexistence through calculation of long-term-growth rates in simulations

Predictions of the central temperature at which *D. pallidifrons* is excluded under variable temperature regimes used the flexible simulation-based method of Ellner *et al*. (2019). This makes asymptotic estimates of the long-term invasion (low-density) population growth rate (IGR) by averaging over long synthetic time series of population dynamics. For each mean temperature, a series of 1000 temperatures were generated: either fixed or with fluctuations (+1.5, 0 or -1.5) following the experimental scenarios.

The population density of the resident *D. pandora* in monoculture was found from simulation using the best-fit population model under both static and variable temperatures regimes (discarding the first 50 steps as a burn-in). The population growth rates from rare of *D. pallidifrons* were calculated for each of these sequences of competitors and temperatures, and the long-term average calculated by the geometric mean. Where for some parameter draw combinations the modelled growth rate was either incalculable (*T* > *T*_*max*_) or below 0.1, a surrogate value of 0.1 was used to facilitate the averaging. These steps were repeated across 1000 draws of the posterior distribution of the model fits and at each central temperature in the range 24-27.6°C in steps of 0.2°C.

Subsequently, to partition out how the environmental fluctuations and competition act together to influence the IGR, further simulation scenarios were explored with and without various processes to isolate the individual and interactive effects. The IGR calculated for the full model (‘Model E’) was partitioned into five components (*r*_0_ + Δ_*C*_ + Δ_*σ*_ + Δ_*_ + Δ_*cov*_) by calculating the differences between models that switched off various processes. To rule out ‘storage-effects’ emerging from covariance between the environmental and competition (Δ_*cov*_), the temperature time series was reshuffled (Model D). To isolate the effect of competition (Δ_*C*_) fluctuations (Δ_*σ*_) and their interaction (Δ_*_), the IGR was calculated with neither competition or fluctuations (Model A), in competition without environmental fluctuations impacting either species (Model B), and as a monoculture with fluctuations (Model C). Table S7 lists the constituent partitions of each model. Note that to cap the total number of partitions, Δ_*_ here incorporates (but is not exactly equivalent to) terms labelled ‘relative-non-linearity’ in similar analyses (e.g. Letten *et al*. 2018).

### Expected time of extinction through direct simulation

To generate a baseline, non-theory derived, forecast for the extinction of *D. pallidifrons*, the selected population models were used in direct simulations of the population dynamics in all four experimental scenarios (with and without competition and environmental variability). Parameter uncertainty was propagated by running the simulations using 1000 distinct posterior draws from both species’ population dynamics models. To assess the role of how stochasticity is modelled, this test was repeated using a model with the same competition kernel, but fit with a Poisson error term.

## Results

### Impact of treatments on time to extinction

Treatment combination had a significant overall effect on the generation at which extirpation was observed (Figure 2, ANOVA: = 8.152, p = 0.0003) but only the slower ‘steady rise monoculture’ treatment was statistically distinct (Tukey HSD test: p< 0.005 for all comparisons with this treatment, p> 0.7 for all others) with a mean expiration 1.73 (SE: 0.358) generations later. When viewed as concurrent stressors, environmental stochasticity and competition had antagonistic interactive effects (Orr *et al*. 2020), in the sense that their combined effect was less than the sum of their individual impacts. The particular trajectory of temperature variability had a significant impact on time-to-extirpation (ANOVA model comparison of Extirpation∼Treatment with Extirpation∼Treatment +(1|Trajectory): = 6.4029, p = 0.011, n = 120, 12 groups). Trajectories with a lower temperature in the first generation were correlated with earlier expiration, but across the 12 trajectories this was not statistically significant (post-hoc linear regression, p = 0.145, = 2.498, adjusted R^2^ = 0.12).

**Figure 2.**
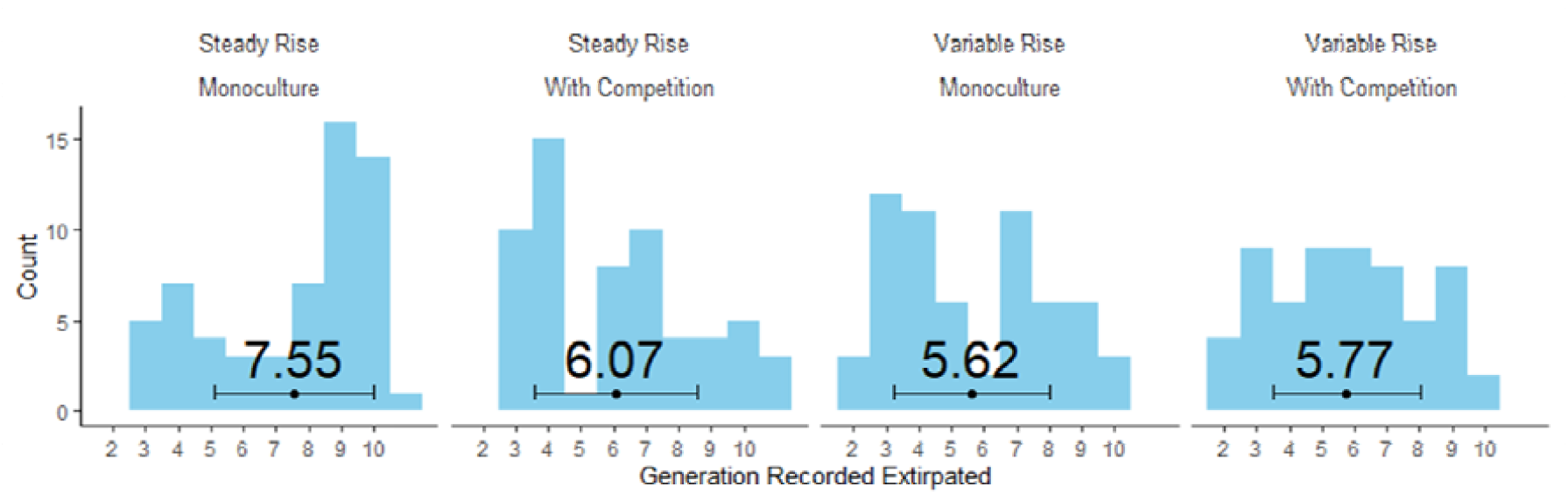
Distribution of observed extirpation points across experimental treatments. Extirpation defined as the first generation where zero *D. pallidifrons* individual were censused. Values shown are means and error bars show +/-1 SD. n=60 per treatment combination (240 total).

### Prediction of time-to-extinction through coexistence theory

With increased temperature, the selected model predicted a shift from *D. pallidifrons* (the resident, highland species) excluding *D. pandora* (the lowland, invader species), through the zone of likely coexistence, to the exclusion of *D. pallidifrons* by *D. pandora* (Figure 3a). Using mean parameter estimates, the threshold for predicted exclusion of *D. pallidifrons* is crossed at approximately 26.7°C, although inspecting the full posterior shows there is support for predicted exclusion somewhat earlier (Figure 3b, red area). Converting these temperatures to generations in the experiment this threshold is first breached in generation 8, just under two generations (corresponding to a 0.8°C difference) after the mean generation of observed extirpation (6.07).

**Figure 3:**
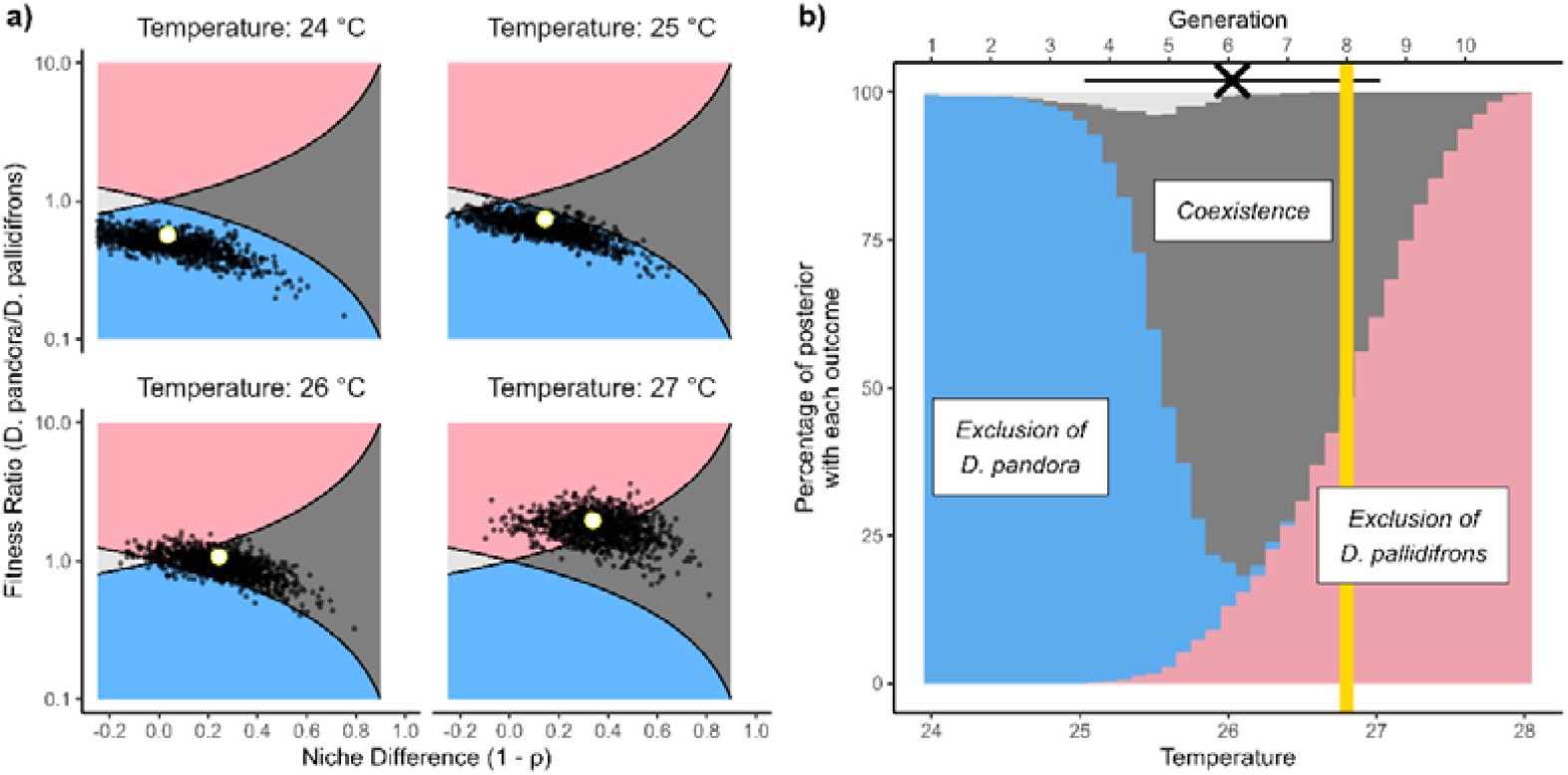
Predictions of coexistence in the steady rise and competition scenario based on analysis of fitted models. a) Full posterior distribution of the niche difference () and fitness differences calculated based on parameter distributions from the selected models. White circles show results based on average parameters. Small black dots show draws from posterior distribution. Shading indicates regions on the plane demarking predicted coexistence outcomes: blue corresponds to predicted exclusion of *D. pandora* at low temperatures, dark grey to the region of predicted coexistence, red to exclusion of *D. pallidifrons* and light grey corresponds to ‘priority effects’, where each species can exclude the other. b) Allocation of posterior support for each of the four possible predicted coexistence outcomes based on inferred niche and fitness differences. Cross at top of figure shows mean observed extirpation (+/-1 SD) of *D. pallidifrons* in the competition and steady rise treatment (see Figure 2 for full distribution).

Modelling the system with simpler models that did not well represent the observed variability gave substantially different predictions. Using a Poisson error distribution coupled with the best fit competition model described above the system is predicted to instead move through a region of priority effects to a sharp shift in predicted dominance of *D. pandora* at an earlier temperature (Figure S6). The use of the same experiments to both fit models from the transitions and test the time of predictions maximises the data available but introduces a certain degree of circularity. Refitting the *D. pallidifrons* model excluding the data from the ‘steady-rise, competition’ treatment used as the main test increases the uncertainty of parameter estimates but gives comparable predictions (posterior support for exclusion at 27.4°C, central 66% interval: 26.5-28.3°C, Figure S7).

Predictions based on the simulation-derived invasion growth rate closely aligned with the analytic prediction (Figure 4a, Model B). The predicted exclusion temperature of *D. pallidifrons* under fluctuating temperatures was 26.23°C (Figure 4a Model E, interpolating linearly between trialled temperature steps), slightly lower than without fluctuations. Predicted IGRs under variable environments show a sharp hinge beyond 26.6°C (Figure 4a). This stemmed from ‘+1.5 °C’ temperatures exceeding causing results past this hinge to be dependent on the arbitrary growth floor. However, the key IGR threshold had already been breached by this point and so this does not impact the key predictions.

**Figure 4.**
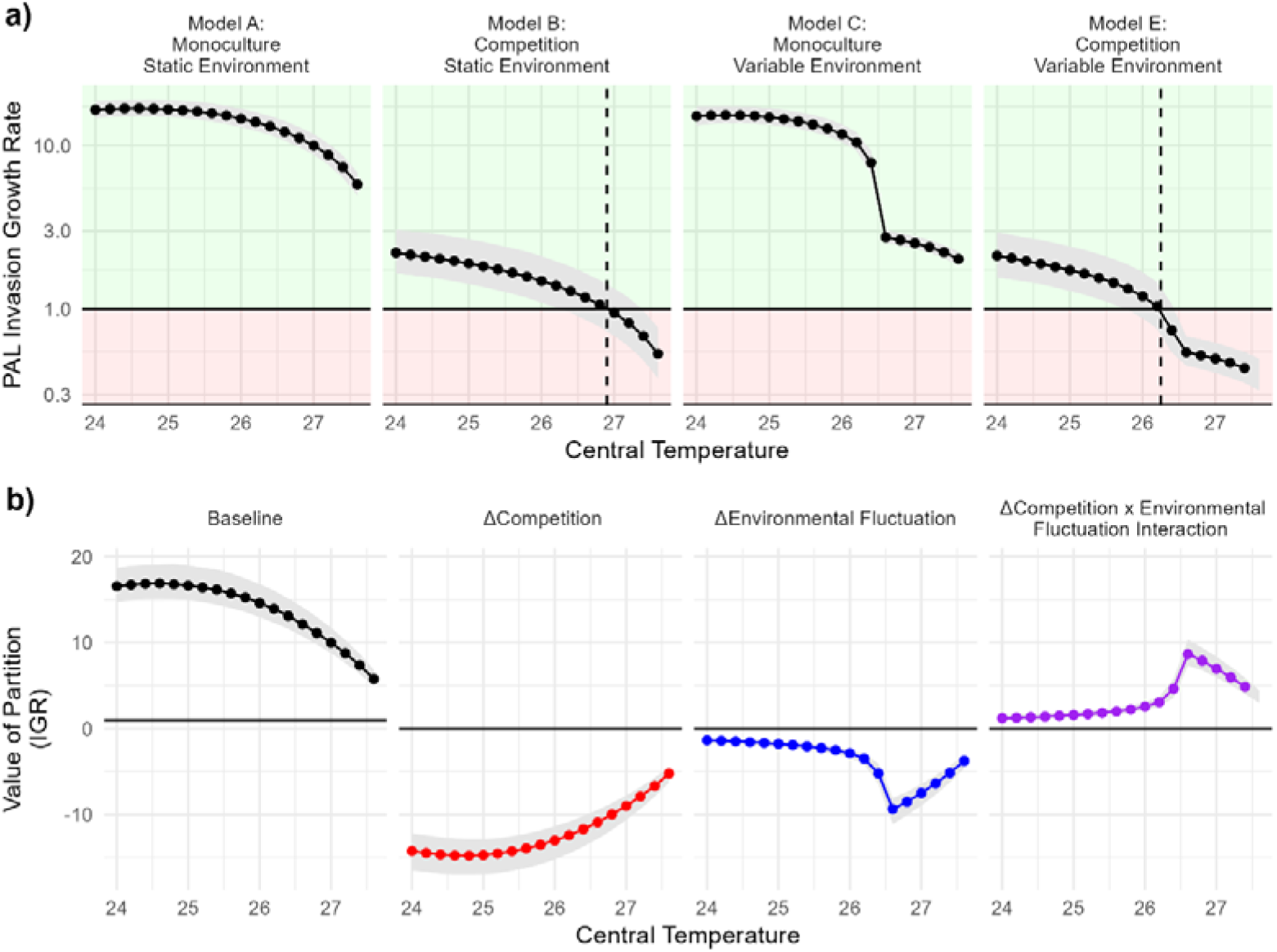
Simulation based decomposition of contributors to the long-term growth rate. Shaded areas show 66% central interval. a) Invasion growth rate (IGR) of *D. pallidifrons* inferred under different simulation scenarios. Results from model D (breaking the covariance between fluctuations in the competitors and the environment) were practically indistinguishable from Model E. b) Value of four key additive partitions from differences in models: *r*_0_, Δ_*C*_, Δ_*σ*_ and Δ_*_(Δ_*COV*_ was negligible).

### Partitioning impact of different processes

The partitioned additive contribution of both competition and temperature fluctuations had negative impacts on the IGR (Figure 4b). At temperatures closer to the transition point the negative pressure from competition reduced while the detrimental impact of the fluctuation increased. Combined, the effects had less joint negative impact – the interaction term between competition and fluctuations was positive, particularly at higher temperatures. As expected from the model structure, there was no detectable ‘storage effect’ component (magnitude of median values of Δ_*cov*_ < 0.0125 for all central temperatures).

### Predictions of extinction by simulation methods

The predictions from the direct simulation of the selected models were generally closer to the observed values than the predictions from coexistence theory (Table 1). However, the direct simulations predict that competition has quite little marginal impact on extinction (hastening extinction by ∼0.6 generations on average), contrary to the observed experimental data and the theory-based approaches. Extinctions from predictions under the simulation model were heavily influenced by the zero-inflated error distribution – simulations derived from a model using Poisson error predicted essentially no extinctions until thermal maximum was reached in G12.

**Table 1.** Predicted generations-until extinction under each experimental scenario. Predictions from the coexistence theory models have been converted from °C to generations for comparability with data. All intervals are central 66%. In the steady rise treatment, simulated extinctions very rarely occurred before the thermal limit causing mean observed generation of extinction to fall outside this central interval.

| Scenario |  | Mean<br>Generation of<br>Observed<br>Extinction | Predictions |  |  |
| --- | --- | --- | --- | --- | --- |
|  |  |  | Coexistence<br>theory | Direct Simulation |  |
| Temperature<br>Trajectory | Competition |  |  | (Zero-Inflated<br>Negative Binomial) | (Poisson) |
| Steady -rise | Competition | 6.07<br>(3.83-9) | 8.04<br>(6.25-9.375) | 6.29<br>(3-9) | 11.90<br>(12-12) |
| Variable-rise | Competition | 5.77<br>(3 -8.17) | 6.57<br>(5.39 - 7.12) | 5.69<br>(3-9) | 10.18<br>(9-12) |
| Steady -rise | Monoculture | 7.55<br>(4 -10) |  | 6.31<br>(3-9) | 11.99<br>(12-12) |
| Variable-rise | Monoculture | 5.62<br>(3-8) |  | 5.73<br>(3-9) | 10.38<br>(9-12) |

## Discussion

In this experimental mesocosm climate-change scenario, coexistence theory was able to make predictions that approximately corresponded with the observations, despite many of the core assumptions not being fulfilled. Although spatial and multi-species effects were excluded by the experimental design, the theory was challenged to deal with non-stationary temporal dynamics, non-finite populations and timeframes, model adequacy as well as potential trait adaptation and positive density dependence. This correspondence appears to also extend to the impact of fluctuating environments, successfully capturing the impact of environmental variability including a strongly antagonistic interaction with competition.

This experiment directly tests coexistence theory both as a predictive tool and as a representative summary of complex ecological dynamics. Nonetheless, assessing the value of the accuracy of the model’s predictions is inevitably somewhat subjective. An optimistic perspective of the alignment would highlight that an estimate within two generations is close enough to be useful. Given the discordance between the theoretical assumptions of modern coexistence theory and the empirical reality of the experiment, it can be seen as surprising and heartening that the results are aligned at all. Further, the experiment does not include any possibility for *D. pallidifrons* recolonising after a stochastic loss, likely speeding the observed extirpation in the mesocosms compared to a more natural situation. A pessimistic view would highlight the substantial temperature difference between observed and predicted extirpations (akin to several decades under most warming scenarios) and the high uncertainty in the prediction, despite levels of replication greatly exceeding the norms of the field. Furthermore, the mean generation of extinction mapped to the temperature with maximum support for coexistence between the species.

Although the full-simulation approach identifies only a small impact of competition on extinction, overall, it is more accurate than the ‘theory-based’ approach. Given then the ready alternative of directly building a simulation model, what can coexistence theory bring to the table? The analytic approach can identify that the combined effect of the warming on multiple temperature dependencies (Figure S4) to cause niche differences to increase (the rightwards drift in Figure 3a). This maintains coexistence longer than may be otherwise expected from an analysis focussing just on population fitness differences. Furthermore, the partitioning process is able to provide further insight on the various mechanisms, isolating a strong antagonistic interactive effect between the two ‘stressors’, that was observed in the experimental data but not predicted in the full simulations. Given the qualitative dependency of the impact of variability on parameters (Terry *et al*. 2022), the successful assessment of the complex impact of variability in this noisy system is notable.

Aligning the scales of experiment and natural scenarios is a particular challenge for extinction dynamics. Experiments with more generations and shallower temperature gradients would allow finer resolution, but will be limited to an even more restricted repertoire of model species. Despite being a controlled mesocosm experiment, individual population trajectories were characterised by a high degree of variation (Figure S1). Counts were complete censuses with populations individually sexed and counted so minimal measurement error can be assumed. Population counts were of the order of dozens (mean of non-zero females counts = 19.9, *σ* =12.7) giving considerable scope for demographic stochasticity (Anderson *et al*. 1982) stemming from individual level-differences within the populations. Within-treatment ‘external’ environmental variability was minimised by the experimental design. Nonetheless, there was likely significant emergent environmental heterogeneity when defined as variation correlated amongst individuals in a vial (Shoemaker *et al*. 2020). Individual vials in the same treatment could be highly divergent, with some vials containing apparently stressed communities with late-developing pupae. Capturing this variation is a major challenge for applying coexistence theory. The best-fit models incorporated zero-inflated negative-binomial error distributions with temperature dependent parameters, a considerably more complex expression than is frequently assumed (Terry & Armitage 2024). Modelling the system with approaches that did not have the flexibility to represent this variability gave different predictions. Although likely chance, it is noteworthy that with a Poisson error model analytic theory gave more accurate and precise predictions of the loss of coexistence (Figure S6) despite direct simulations with Poisson error model being highly inaccurate.

It must nonetheless be acknowledged that the alignments observed in this experiment under controlled conditions likely represent a ‘best-case’ expectation of accuracy. Beyond the model and parameter uncertainty captured within ecological modelling, natural world forecasts must also grapple with climate change uncertainty (incorporating both human and earth system responses, Beaumont *et al*. 2008). The extent and accuracy of data available in this test likely exceeds that available for many applied cases. While the forecasts could be considered acceptably accurate on average, the high heterogeneity among replicates suggests strong fundamental limits of predictability at local scales. This heterogeneity arises even without incorporating dispersal variation (Melbourne & Hastings 2009) or the true field community complexity, including more competitive rivals and strong interactions with parasitoid natural enemies (Chen & Lewis 2023). Rigorously applying coexistence theory to more complex communities requires significant extensions to the analysis approach to tackle alternative invasion pathways (Chesson 2018; Ranjan *et al*. 2024; Spaak & Schreiber 2023), which are likely to place unrealistic demands on model parameterisation.

Strong positive density dependence can challenge the calculation and use of an invasion growth rate and are not represented in the fitted Beverton-Holt model. Impactful Allee effects, where solitary mated females reduce the rate of eggs they lay, have been observed with these species in previous competition experiments (Terry *et al*. 2021). In *D. melanogaster* this social modulation effect has been well studied and is possibly associated with the need to overcome competition with fungi (Bailly *et al*. 2023; Wertheim *et al*. 2002). Nonetheless, in this experiment there were few cases with only one or two females and Allee effects appear to have had little impact on the dynamics.

The modelling framework used here assumes fixed traits, building in neither potential cross-generational acclimatory responses or genotypic adaptation. Acclimatation effects across and within generations in *Drosophila* can be considerable (Cavieres *et al*. 2020; Schiffer *et al*. 2013) and rapid evolutionary changes are also possible (e.g. Rudman *et al*. 2022), although the high intrinsic variation and relatively restricted range of temperatures may be expected to lead to comparatively little evolutionary genetic change over these timescales. While several authors have proposed extensions that incorporate evolutionary feedbacks (Pastore *et al*. 2021; Yamamichi *et al*. 2022), the high parametrisation requirements mean it is rarely incorporated into empirical applications. As such, for the purposes of this study ‘no-trait changes’ is effectively one of the assumptions under investigation. The temperature trajectory in this experiment will lead to a close match between temperatures across transitions, and as such any effects of acclimation are likely to be ‘built-in’ to the fitted thermal impacts.

In natural systems, the *Drosophila* exist in a highly dynamic metacommunity as individual ephemeral food resources become available and are consumed. Such a spatially distributed system will likely mitigate the substantial failure rate of individual patches. Further, within the mesocosms there was no distinction between the conditions under which the species’ performance was assessed and those for the long-term experiment. In field conditions it can be hard to capture environmental impacts on all life-stages appropriately, and indeed each may undergo distinct environmental trajectories (Sinclair *et al*. 2016). Nonetheless, this study offers support that when carefully applied even quite abstract coexistence models can indeed make informative forecasts for contemporary change beyond the precise bounds of their formal applicability.

As communities reorganise in the face of changing environments, trustworthy tools are needed to offer guidance. There has been substantial and ongoing debate about how best to address the gap between the asymptotic theoretical world and the finite and messy reality that we must confront (Pande *et al*. 2020; Schreiber *et al*. 2023). Ultimately, observation and experimentation are essential to offer direction. Since there is not the luxury of sufficient time to conduct equivalent direct validation experiments in species with longer lifecycles, mesocosm experiments with high-turnover model species will have a major role to play. Validation tests such as those presented here can contextualise the expected accuracy from models to develop an expectation of the appropriate level of confidence in, and hence application and deployment of, coexistence theory.

## Acknowledgments

JCDT was funded by a Leverhulme Trust Early Career Fellowship (ECF-2022-666). The help and support from the Oxford Biology Fly Lab and Community Ecology research groups is gratefully acknowledged. Highly constructive comments from three anonymous reviewers substantially improved the manuscript.

## Notes

https://doi.org/10.5281/zenodo.10679556

### Competing Interest Statement

The authors have declared no competing interest.

### Summary of Updates

Writing tweaks, updates to references, correction to Table 1.

