## Supplementary Information for "An experimental validation test of ecological coexistence theory to forecast extinction under rising temperatures"

J. Christopher D. Terry^1*^

1: Department of Biology, University of Oxford, 11A Mansfield Road, Oxford, OX1 3SZ

 0000-0002-0626-9938

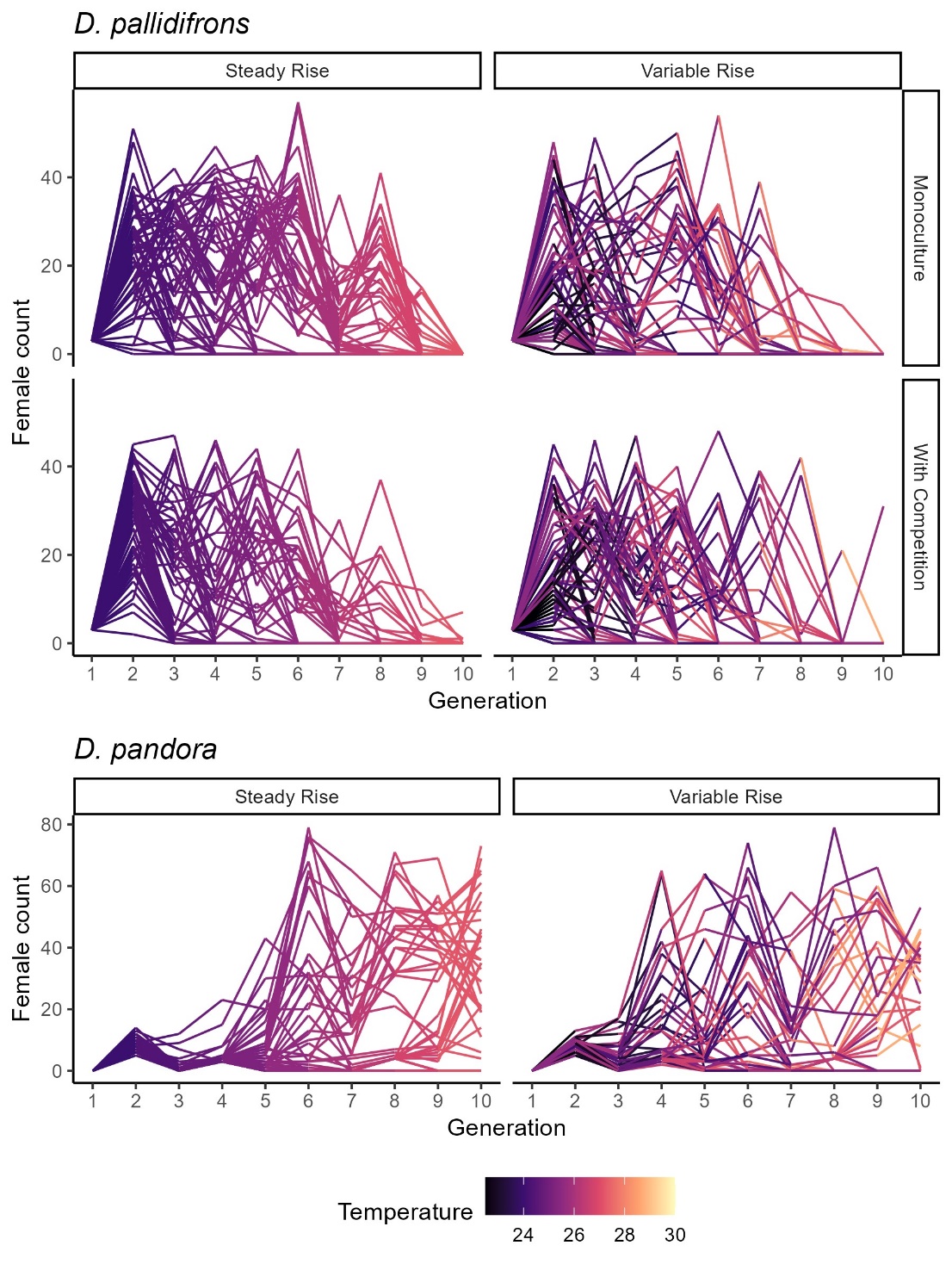

**Figure S1** Raw time series of female counts for each population line during the experiment facetted by treatment and coloured by incubator temperature.

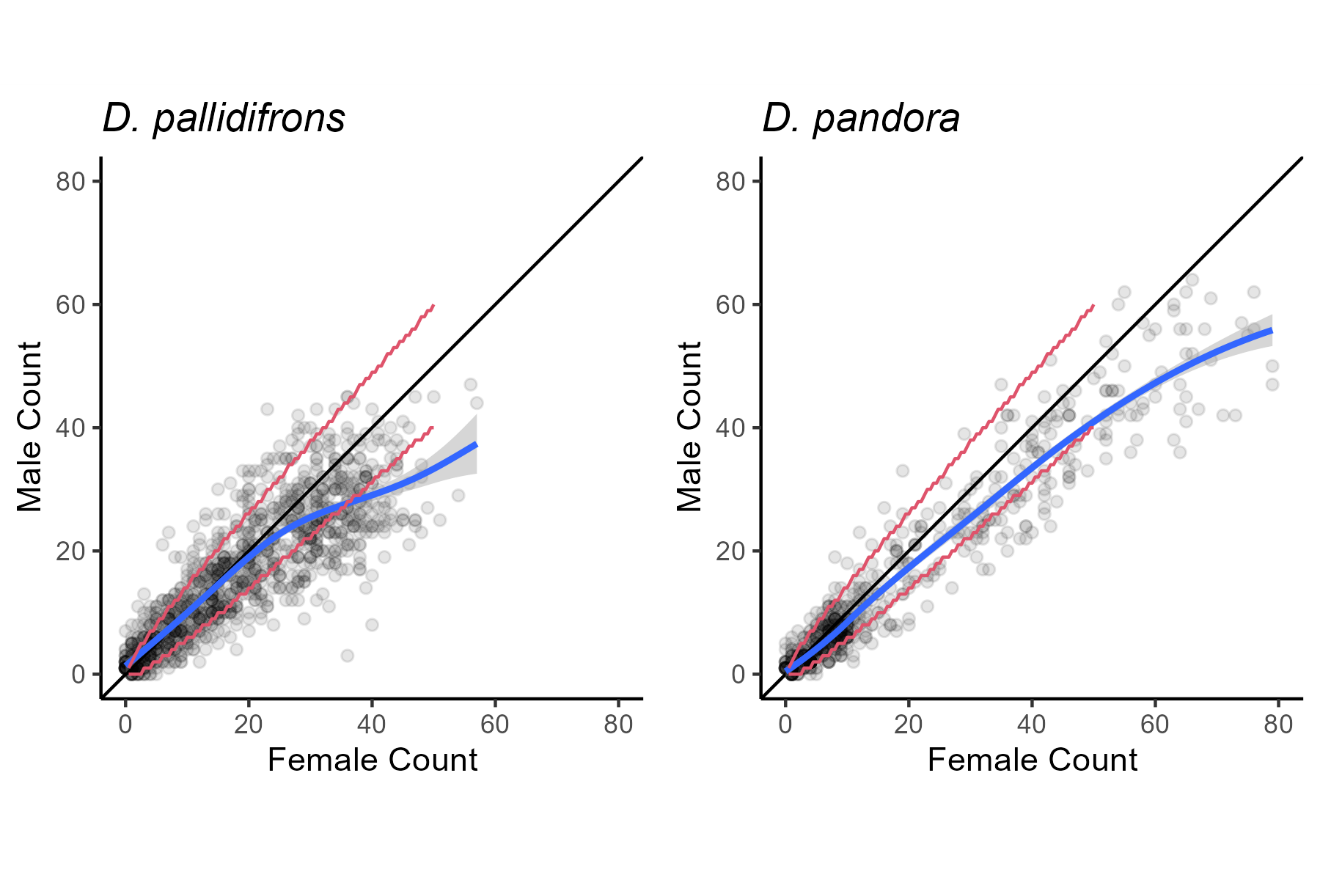
**Figure S2** Sex ratio of the two species with respect to density. Red lines show 95% interval of assuming a 1:1 ratio. Blue line shows a smooth loess plot highlighting female skew at high overall densities.

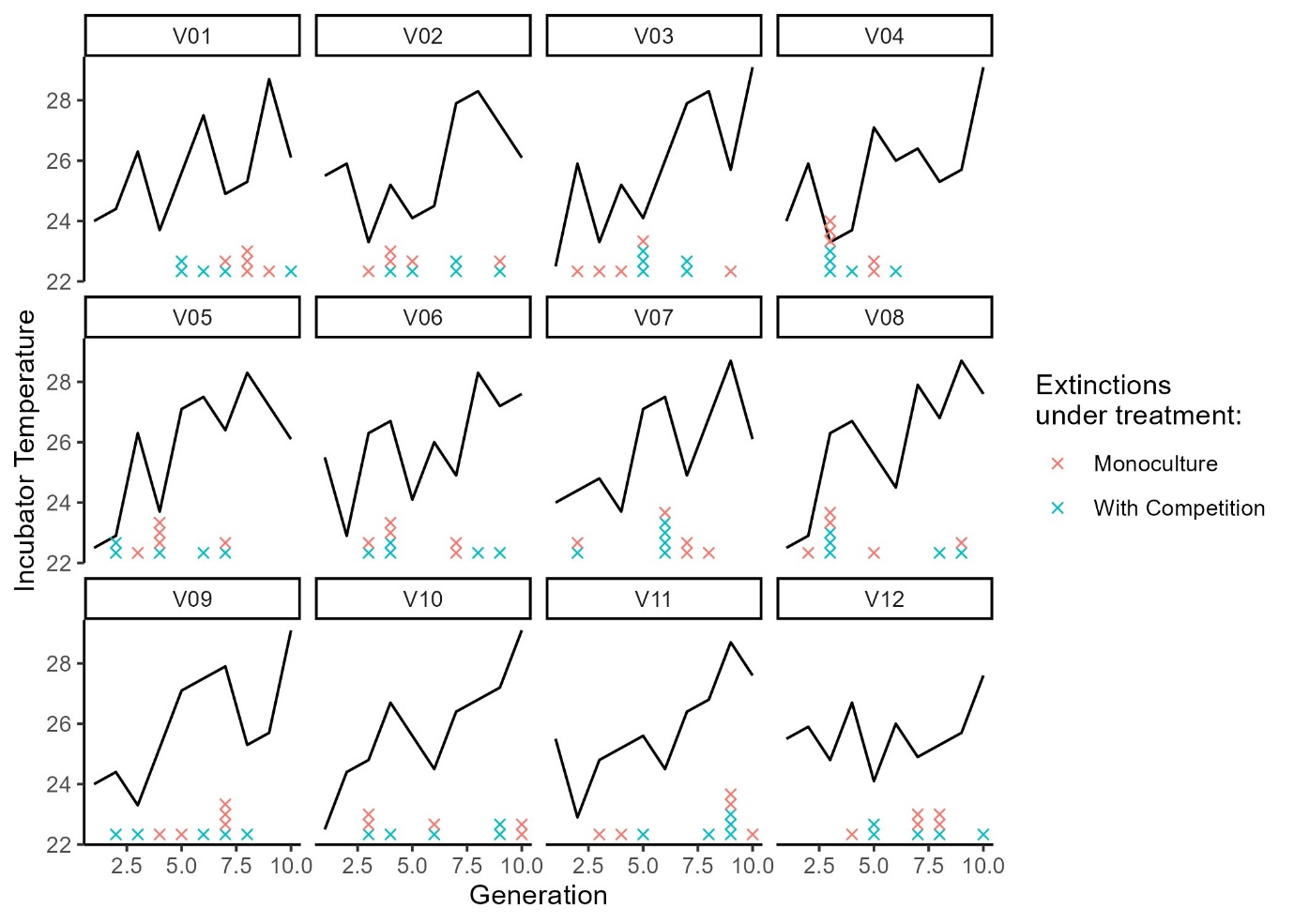

**Figure S3.** Extinctions within each variability trajectory within the ‘Variable’ temperature treatment.

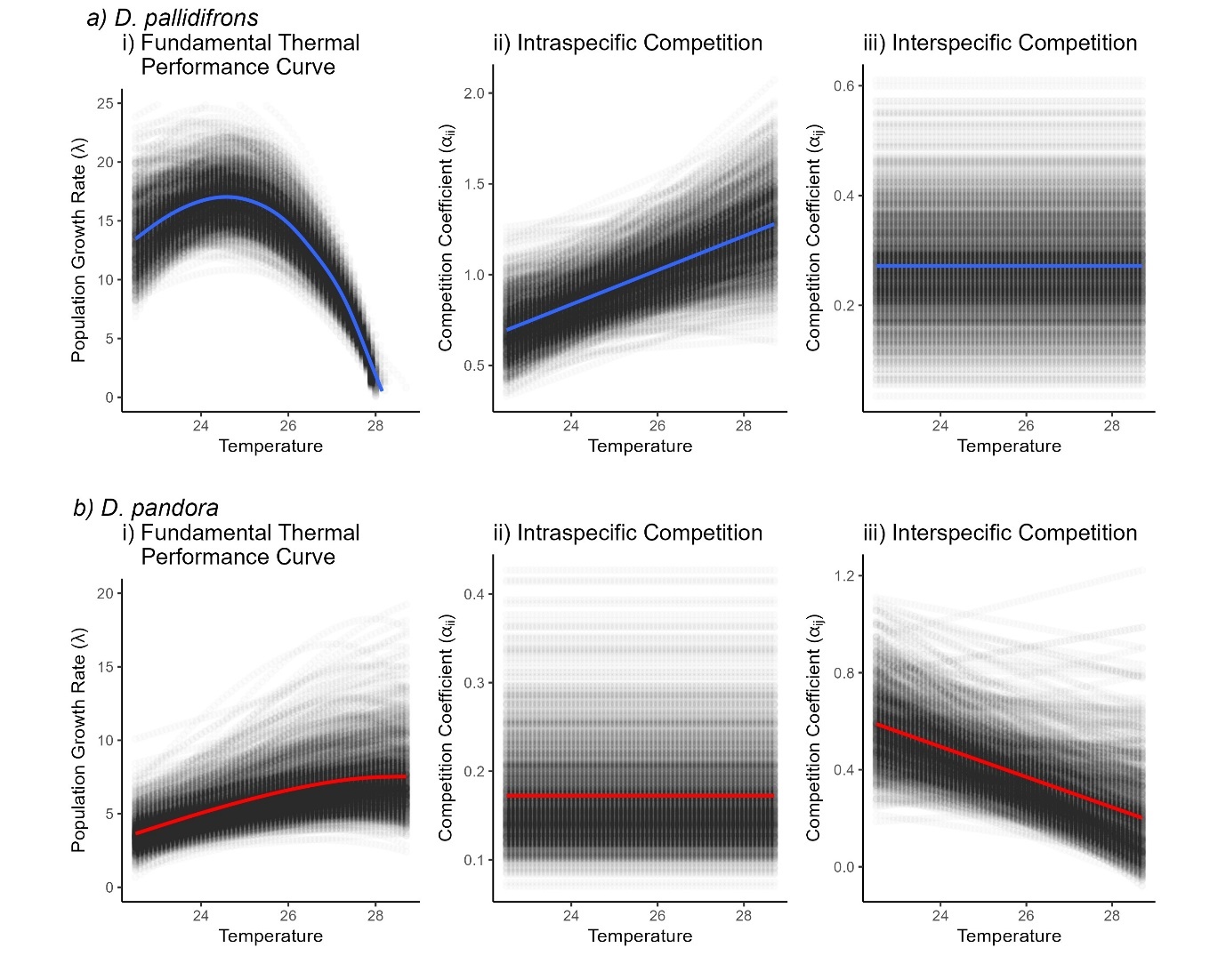

**Figure S4** Posterior distribution of dependence of composite parameters ($\lambda$ and $\alpha$ competition coefficients) on temperature for the selected model for each species. Coloured lines show a smooth fit through the posterior distribution.

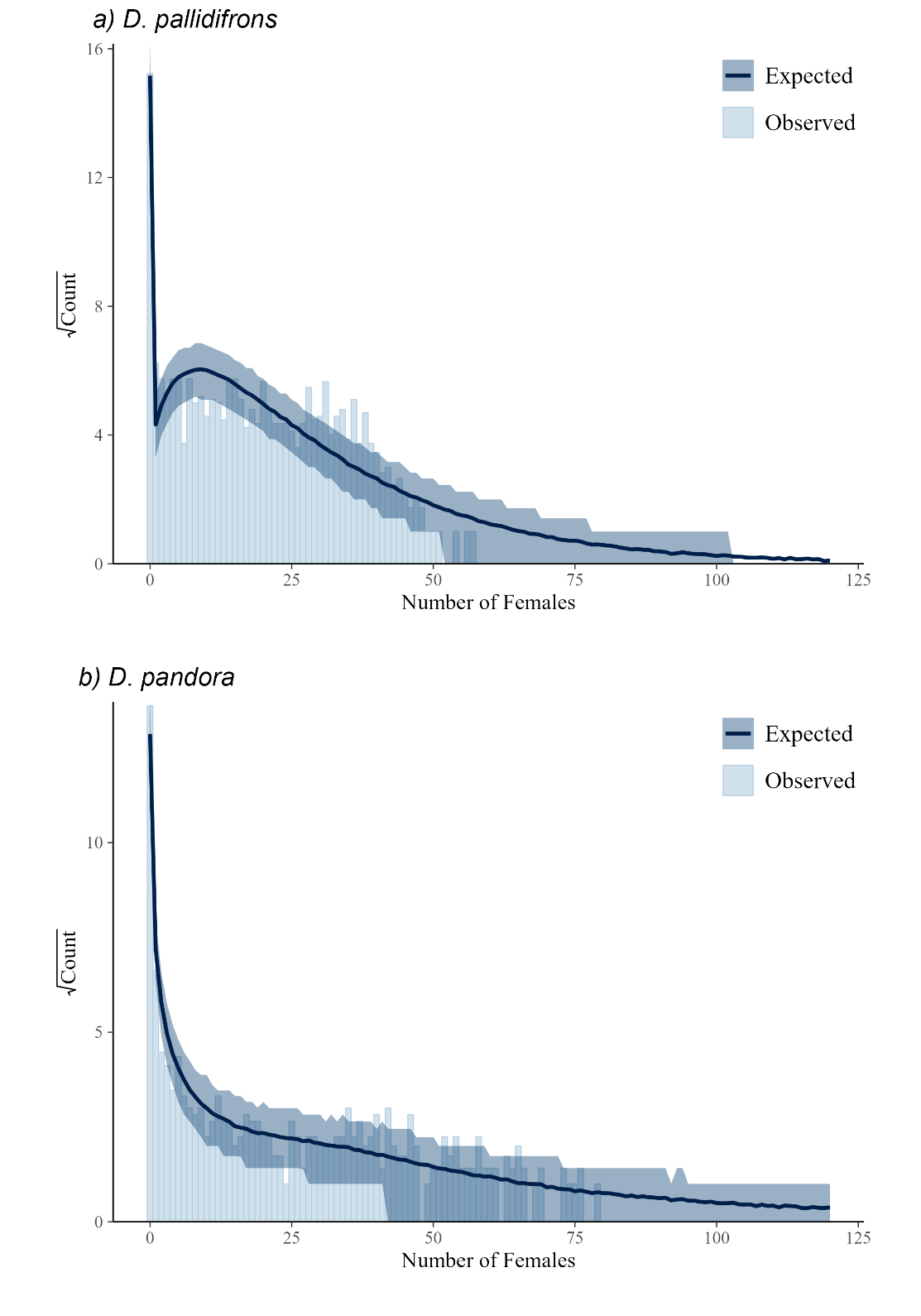

**Figure S5.** Posterior predictive distribution of selected model through rootograms^1^**.** Overall, the modes can perform well at low levels, including the zero-inflation, although the model predicts more high counts of both species than are observed.

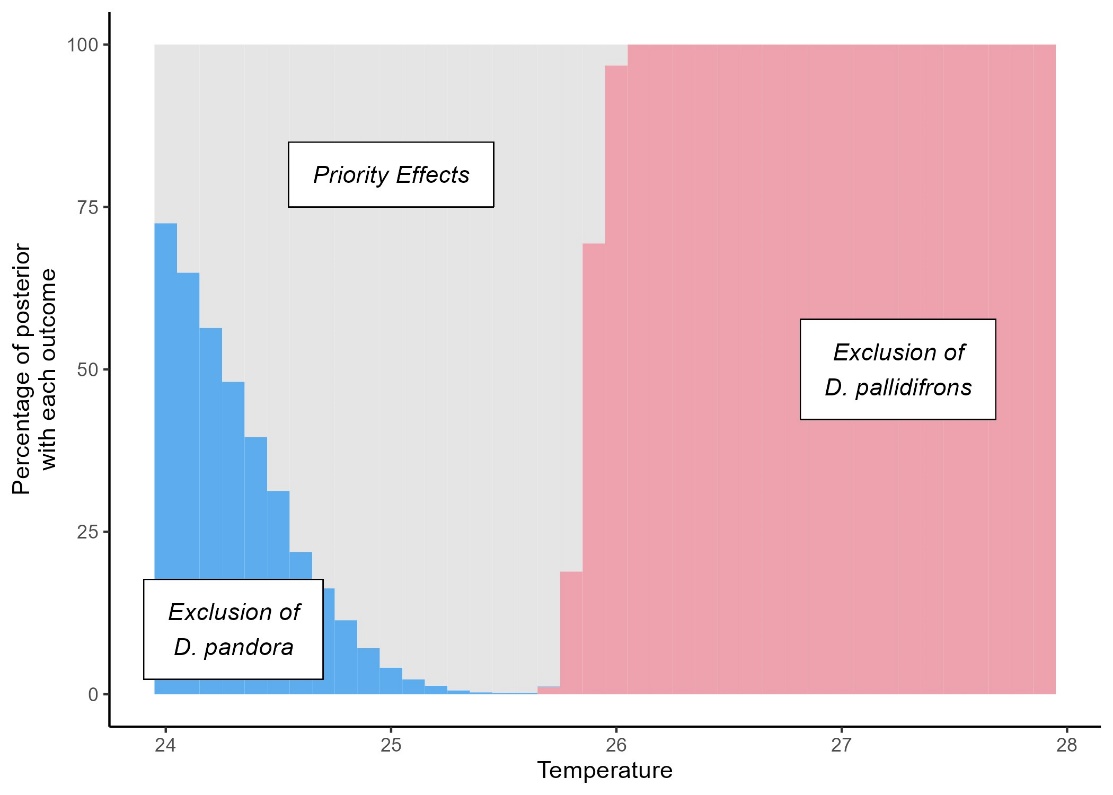

**Figure S6.** Coexistence predictions using the same model competition and thermal performance model structure, but a Poisson error distribution. Compared to the principal model, this alternative identifies a period of priority effects rather than coexistence and a much sharper transition. Despite poorly capturing the observed error distribution, it is notable that it gives both an accurate and precise prediction of the point of exclusion.

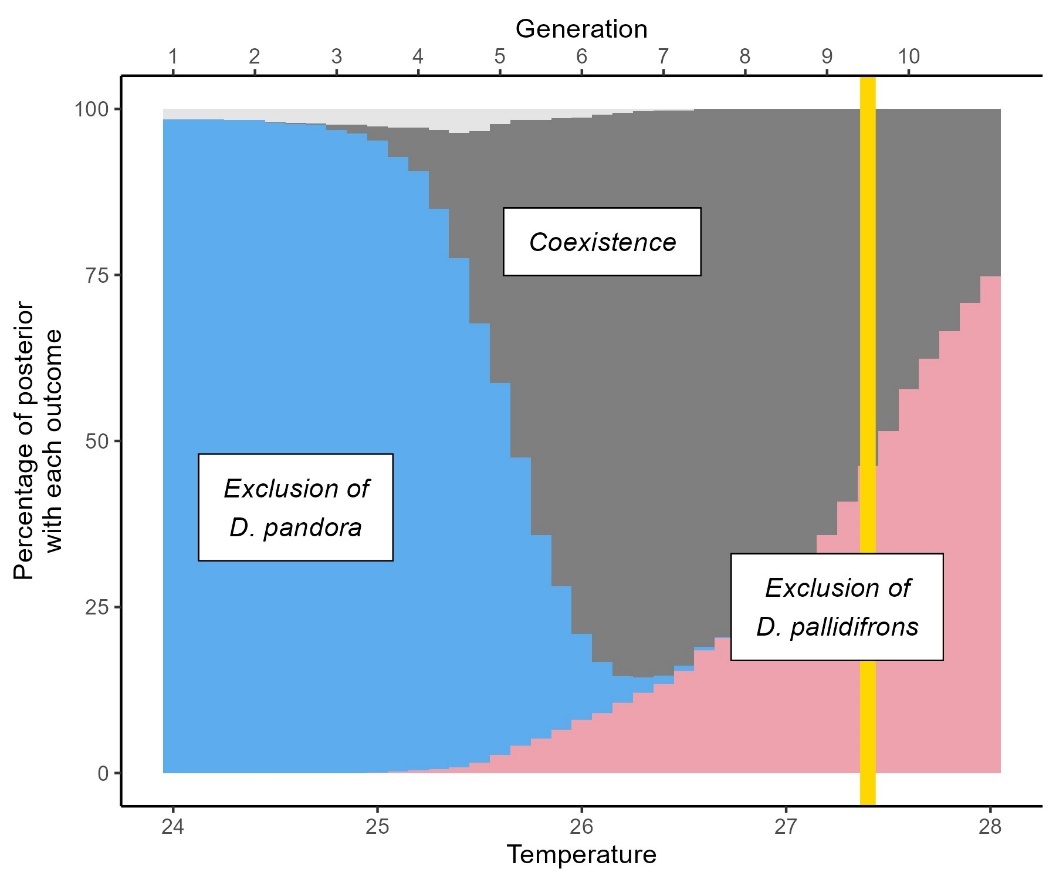

**Figure S7** Coexistence predictions where *D. pallidifrons* model is trained without transition data from the ‘steady rise, competition’ experimental treatment used as the principle test.

**Table S1.** Tested competition model structures and dependencies. To prevent combinatorial explosion, models chosen to be tested were developed through sequential modification of initial candidate models based on observed fitted parameters.

| **Model Name**  **(BH = Beverton-Holt)** | **Model formula for competition kernel** | **N** |
| --- | --- | --- |
| Lotka-Volterra (excluded due to poor convergence) | $1+{\alpha_{i,PAL}N}_{PAL,t}+\alpha_{i,PAN}N_{\mathrm{PAN},t}$ | **2** |
| BH | $\frac{1}{1+{\alpha_{i,PAL}N}_{PAL,t}+\alpha_{i,PAN}N_{\mathrm{PAN},t}}$ | **2** |
| Beta-BH | $\frac{\boldsymbol{1}}{\left( 1+{\alpha_{i,PAL}N}_{PAL,t}+\alpha_{i,PAN}N_{\mathrm{PAN},t} \right)^{\boldsymbol{\beta}}}$ | **3** |
| Two-way Temperature dependent BH | $\frac{\boldsymbol{1}}{1+\left( \alpha_{0i,PAL}+\alpha_{Ti,PAL}T_{t} \right)N_{PAL,t}+\left( \alpha_{0i,PAN}+\alpha_{Ti,PAN}T_{t} \right)N_{PAN,t}}$ | **4** |
| Two-way Temperature dependent Beta- BH | $\frac{\boldsymbol{1}}{\left( 1+\left( \alpha_{0i,PAL}+\alpha_{Ti,PAL}T_{t} \right)N_{PAL,t}+\left( \alpha_{0i,PAN}+\alpha_{Ti,PAN}T_{t} \right)N_{PAN,t} \right)^{\boldsymbol{\beta}}}$ | **5** |
| PAL-only Temperature dependent BH | $\frac{1}{1+\left( \alpha_{0i,PAL}+\alpha_{Ti,PAL}T_{t} \right)N_{PAL,t}+\alpha_{i,PAN}N_{j,t}}$ | **3** |
| PAL-only Temperature dependent Beta- BH | $\frac{\boldsymbol{1}}{\left( 1+\left( \alpha_{0i,PAL}+\alpha_{Ti,PAL}T_{t} \right)N_{PAL,t}+\alpha_{i,PAN}N_{PAN,t} \right)^{\boldsymbol{\beta}}}$ | **4** |
| PAL-only Temperature dependent BH with log-competition | $\frac{1}{1+\left( \alpha_{0i,PAL}+\alpha_{Ti,PAL}T_{t} \right){\log(N}_{PAL,t}+1)+\alpha_{i,PAN} log(N_{PAN,t}+1)}$ | **3** |
| PAL-only Temperature dependent Beta-BH with log-competition | $\frac{\boldsymbol{1}}{\left( 1+\left( \alpha_{0i,PAL}+\alpha_{Ti,PAL}T_{t} \right){\log(N}_{PAL,t}+1)+\alpha_{i,PAN}log(N_{PAN,t}+1) \right)^{\boldsymbol{\beta}}}$ | **4** |

**Table S2.** Tested error model structures and dependencies with *brms* package R code syntax for additional clarity. This uses the negative binomial parameterisation in terms of mean (or location, $\mu$) and a shape parameter ($\phi$) such that $E\left[ n \right]=\mu\mathrm{and} Var\left[ n \right]=\mu+\frac{\mu^{2}}{\phi}$.

| **Model Name**  **(NB = Negative-Binomial)** | **Code used within *brms*** |
| --- | --- |
| Poisson | family = poisson(link = "identity") |
| Negative Binomial | family = negbinomial(link = "identity") |
| Zero-inflated NB | family = zero_inflated_negbinomial(link = "identity") |
| Temperature-dependent-zero-inflated NB | family = zero_inflated_negbinomial(link = "identity")  additional terms: zi~PrevTemp |
| Zero-inflated NB with competition dependent shape | family = zero_inflated_negbinomial(link = "identity")  additional terms: shape~Prev_PAL_FEMALE or shape~Prev_PAN_FEMALE |
| Zero-inflated NB with temperature dependent shape | family = zero_inflated_negbinomial(link = "identity")  additional terms: shape~PrevTemp |

**Table S3.** Relative performance of the top 25 models (out of 48 that were comparable) to fit the *D. pallidifrons* transition data. All tested models used the Briere thermal performance curve. R-hat convergence is of the most poorly converged parameter in the model.

| Competition function | Error Model | $\Delta$ELPD LOO | R-hat Convergence |
| --- | --- | --- | --- |
| BHTEMP1Beta | Temperature-dependent-zero-inflated NB | 0 | 1.001 |
| BHTEMPBeta | Temperature-dependent-zero-inflated NB | -0.308 | 1.003 |
| BHTEMP | Temperature-dependent-zero-inflated NB | -6.1 | 1.002 |
| BHTEMP1 (x) | Temperature-dependent-zero-inflated NB | -6.29 | 1.002 |
| BHBeta | Temperature-dependent-zero-inflated NB | -7.42 | 1.002 |
| BH | Temperature-dependent-zero-inflated NB | -9.14 | 1.002 |
| BHTEMPBeta | Zero-inflated NB with temperature-dependent shape | -11.7 | 1.014 |
| BHTEMP1LogBeta | Temperature-dependent-zero-inflated NB | -13.4 | 1.002 |
| BHTEMP | Zero-inflated NB with temperature-dependent shape | -19.4 | 1.002 |
| BHTEMP1 | Zero-inflated NB with temperature-dependent shape | -20.4 | 1.001 |
| BHTEMP1LogBeta | Zero-inflated NB with temperature-dependent shape | -25.5 | 1.003 |
| BHBeta | Zero-inflated NB with temperature-dependent shape | -25.7 | 1.001 |
| BH | Zero-inflated NB with temperature-dependent shape | -28.4 | 1.001 |
| BHTEMPBeta | Zero-inflated NB | -37 | 1.001 |
| BHTEMP1Beta | Zero-inflated NB | -37.1 | 1.003 |
| BHTEMPBeta | Zero-inflated NB with competition-dependent shape | -37.7 | 1.002 |
| BHTEMP1Beta | Zero-inflated NB with competition-dependent shape | -38.3 | 1.003 |
| BHTEMP | Zero-inflated NB | -42.3 | 1.003 |
| BHBeta | Zero-inflated NB | -42.5 | 1.001 |
| BHTEMP1 | Zero-inflated NB | -42.6 | 1.002 |
| BHTEMP | Zero-inflated NB with competition-dependent shape | -43.5 | 1.004 |
| BHBeta | Zero-inflated NB with competition-dependent shape | -43.7 | 1.002 |
| BHTEMP1 | Zero-inflated NB with competition-dependent shape | -43.7 | 1.002 |
| BH | Zero-inflated NB | -44.2 | 1.002 |
| BH | Zero-inflated NB with competition-dependent shape | -45.5 | 1.002 |

**Table S4** Relative performance of the top 25 models (out of 48 that were comparable) to fit the *D. pandora* transition data. All tested models used the Briere thermal performance curve. R-hat convergence is of the most poorly converged parameter in the model.

| Competition function | Error Model | $\Delta$ELPD LOO | R-hat Convergence |
| --- | --- | --- | --- |
| BHTEMP1 | Zero-inflated NB with competition-dependent shape | 0 | 1.002 |
| BHTEMP | Zero-inflated NB with competition-dependent shape | -0.53 | 1.002 |
| BHTEMP1Beta | Zero-inflated NB with competition-dependent shape | -1.29 | 1.005 |
| BHTEMPBeta | Zero-inflated NB with competition-dependent shape | -1.58 | 1.01 |
| BH | Zero-inflated NB with competition-dependent shape | -2.15 | 1.002 |
| BHBeta | Zero-inflated NB with competition-dependent shape | -2.72 | 1.002 |
| BHTEMP1LogBeta | Zero-inflated NB with competition-dependent shape | -12.8 | 1.002 |
| BHTEMP1 | Zero-inflated NB with temperature-dependent shape | -26.1 | 1.001 |
| BHTEMP | Zero-inflated NB with temperature-dependent shape | -26.3 | 1.001 |
| BHTEMPBeta | Zero-inflated NB with temperature-dependent shape | -27.7 | 1.003 |
| BHTEMP1Beta | Zero-inflated NB with temperature-dependent shape | -27.9 | 1.003 |
| BH | Zero-inflated NB with temperature-dependent shape | -31.3 | 1.002 |
| BHBeta | Zero-inflated NB with temperature-dependent shape | -31.8 | 1.002 |
| BHTEMP1LogBeta | Zero-inflated NB with temperature-dependent shape | -34.5 | 1.001 |
| BHTEMP1Log | Zero-inflated NB with competition-dependent shape | -39.6 | 1.001 |
| BH | Temperature-dependent-zero-inflated NB | -50.9 | 1.002 |
| BHTEMP1 | Temperature-dependent-zero-inflated NB | -50.9 | 1.002 |
| BHTEMP | Temperature-dependent-zero-inflated NB | -51.4 | 1.003 |
| BHBeta | Temperature-dependent-zero-inflated NB | -51.4 | 1.003 |
| BHTEMP1Beta | Temperature-dependent-zero-inflated NB | -51.9 | 1.002 |
| BHTEMPBeta | Temperature-dependent-zero-inflated NB | -52 | 1.002 |
| BHTEMP1LogBeta | Temperature-dependent-zero-inflated NB | -59.6 | 1.001 |
| BHTEMP1Log | Zero-inflated NB with temperature-dependent shape | -64 | 1.001 |
| BHTEMP1 | Zero-inflated NB | -67.1 | 1.001 |
| BHTEMP | Zero-inflated NB | -67.9 | 1.002 |

**Table S5** Posterior summary of raw parameters within selected model for *D. pallidifrons*. Model followed: PAL_FEMALE ~ Prev_PAL_FEMALE * (B0 * PrevTemp * (PrevTemp - Tmin) * sqrt(Tmax - PrevTemp)) * (1/((1 + (alphaii0 + alphaiiT * Temp20) * Prev_PAL_FEMALE_comp + alphaij * Prev_PAN_FEMALE))) , zi ~ PrevTemp See code supplement for full model code and full posterior distributions. Note that to aid readability and make it easier to define weak priors on the intercept, the temperature value used to determine the effect of temperature on competition impacts was reduced by 20 to create the ‘Temp20’ variable.

| Parameter | Estimate | Est.Error | l-95% CI | u-95% CI |
| --- | --- | --- | --- | --- |
| B0 | 0.0795 | 0.0257 | 0.0332 | 0.13 |
| Tmin | 18.6 | 2.18 | 12.4 | 21 |
| Tmax | 28 | 0.0605 | 27.9 | 28.1 |
| alphaii0 | 0.456 | 0.226 | 0.0555 | 0.921 |
| alphaij | 0.273 | 0.0954 | 0.106 | 0.478 |
| alphaiiT | 0.0955 | 0.0421 | 0.015 | 0.175 |
| Zi_Intercept | -17.4 | 2.02 | -21.4 | -13.6 |
| zi_PrevTemp | 0.628 | 0.0788 | 0.48 | 0.784 |

**Table S6** Posterior summary of raw parameters within selected model for *D. pandora*. Where model followed: PAN_FEMALE ~ Prev_PAN_FEMALE * (B0 * PrevTemp * (PrevTemp - Tmin) * sqrt(Tmax - PrevTemp)) * (1/((1 + alphaii * Prev_PAN_FEMALE_comp + (alphaij0 + alphaijT * Temp20) * Prev_PAL_FEMALE))) , shape ~ Prev_PAN_FEMALE See code supplement for full model code and full posterior distributions. Note that to aid readability and make it easier to define weak priors on the intercept, the temperature value used to determine the effect of temperature on competition impacts was reduced by 20 to create the ‘Temp20’ variable.

| Parameter | Estimate | Est.Error | l-95% CI | u-95% CI |
| --- | --- | --- | --- | --- |
| B0 | 0.0149 | 0.00929 | 0.0043 | 0.0393 |
| Tmin | 17.8 | 2.73 | 11.2 | 21.5 |
| Tmax | 33.6 | 2.99 | 29.5 | 39.5 |
| alphaii | 0.173 | 0.0566 | 0.0974 | 0.314 |
| alphaij0 | 0.736 | 0.22 | 0.341 | 1.21 |
| alphaijT | -0.0607 | 0.0336 | -0.123 | 0.00964 |
| Shape_intercept | -1.14 | 0.114 | -1.36 | -0.903 |
| shape_Prev_PAN_FEMALE | 0.0505 | 0.00403 | 0.0424 | 0.0583 |

| **Model** | **Competition from PAN** | **Environment** | **Constituent Partitions** |
| --- | --- | --- | --- |
| A (Baseline) | None | Fixed | $r_{0}$ |
| B | Constant (carrying capacity at constant environment) | Fixed | $r_{0}+\Delta_{C}$ |
| C | None | Fluctuating | $r_{0}+\Delta_{\sigma}$ |
| D | Present (variable following fluctuations) | Fluctuating (uncorrelated between focal and competitor) | $r_{0}+\Delta_{C}+\Delta_{\sigma}+\Delta_{*}$ |
| E (Full) | Present (variable following fluctuations) | Fluctuating (Same sequence affects both focal and competitor) | $r_{0}+\Delta_{C}+\Delta_{\sigma}+\Delta_{*}+\Delta_{cov}$ |

**Table S7** Models compared to partition contribution of processes to changes in the predicted long-term growth rate. Care must be taken when comparing between studies as different choices in how the environmental variation impacts the competitor species density can create additional small interactive effects.
